# Genome Wide Association Analyses Based on Broadly Different Specifications for Prior Distributions, Genomic Windows, and Estimation Methods

**DOI:** 10.1101/120808

**Authors:** Chunyu Chen, Juan P. Steibel, Robert J. Tempelman

## Abstract

A popular strategy (EMMAX) for genome wide association (GWA) analysis fits all marker effects as classical random effects (i.e., Gaussian prior) by which association for the specific marker of interest is inferred by treating its effect as fixed. It seems more statistically coherent to specify all markers as sharing the same prior distribution, whether it is Gaussian, heavy-tailed (BayesA), or has variable selection specifications based on a mixture of, say, two Gaussian distributions (SSVS). Furthermore, all such GWA inference should be formally based on posterior probabilities or test statistics as we present here, rather than merely being based on point estimates. We compared these three broad categories of priors within a simulation study to investigate the effects of different degrees of skewness for quantitative trait loci (QTL) effects and numbers of QTL using 43,266 SNP marker genotypes from 922 Duroc-Pietrain F2 cross pigs. Genomic regions were based either on single SNP associations, on non-overlapping windows of various fixed sizes (0.5 to 3 Mb) or on adaptively determined windows that cluster the genome into blocks based on linkage disequilibrium (LD). We found that SSVS and BayesA lead to the best receiver operating curve properties in almost all cases. We also evaluated approximate marginal a posteriori (MAP) approaches to BayesA and SSVS as potential computationally feasible alternatives; however, MAP inferences were not promising, particularly due to their sensitivity to starting values. We determined that it is advantageous to use variable selection specifications based on adaptively constructed genomic window lengths for GWA studies.

**SUMMARY:** Genome wide association (GWA) analyses strategies have been improved by simultaneously fitting all marker effects when inferring upon any single marker effect, with the most popular distributional assumption being normality. Using data generated from 43,266 genotypes on 922 Duroc-Pietrain F2 cross pigs, we demonstrate that GWA studies could particularly benefit from more flexible heavy-tailed or variable selection distributional assumptions. Furthermore, these associations should not just be based on single markers or even genomic windows of markers of fixed physical distances (0.5 − 3.0 Mb) but based on adaptively determined genomic windows using linkage disequilibrium information.

## INTRODUCTION

Recent developments in genotyping technology have made single nucleotide polymorphism (SNP) genotype marker panels, based on thousands, and now millions, of markers, available for many livestock species (Wiggans *et al*. 2013; Kemper *et al*. 2015). Genome wide association (**GWA**) analyses have been increasingly used to help pinpoint regions containing potential causal variants or quantitative trait loci (**QTL**) for economically important phenotypes based on fitting SNP markers as covariates. An increasingly popular inferential approach for GWA is based on fitting phenotypes as a joint linear function of all markers using mixed-model procedures such as those invoked in the popular EMMAX procedure (Kang *et al*. 2010) and other similar procedures (Lippert *et al*. 2011; Zhou and Stephens 2012). Jointly accounting for all SNP effects when inferring upon a specific SNP marker of interest generally improves precision and power while also accounting for potential population structure (Kang *et al*. 2008).

Now GWA inferences in EMMAX and related procedures are based on treating the effect of the SNP marker of interest as fixed with all other marker effects as normally distributed random effects, noting that this process is repeated in turn for every single marker. These “fixed effects” hypothesis tests are based on generalized least squares (GLS) inference, with P-values being subsequently adjusted for the total number of markers or tests. Goddard *et al*. (2016) have recently pointed out the paradox with of treating markers as fixed for inference but then otherwise as random to account for population structure for inference on association with other markers. Random effects modeling with all SNP effects treated as random, including the one of inferential interest, is synonymous with shrinkage based inference. Shrinkage or posterior inference has been demonstrated to facilitate reliable inference without any formal requirements for multiple comparison adjustments (Gelman *et al*. 2012). However, with SNP markers treated as identically and independently distributed variables from a Gaussian distribution, the resulting shrinkage from random effects modeling can be too “hard”, particularly with greater marker densities (Hayes 2013). Subsequently, this random effects test has been deemed to be far too conservative in various applications, as further demonstrated by Gualdron Duarte *et al*. (2014).

Prior specifications that are sparser than Gaussian may be more important for GWA since they more likely better characterize the true genetic architecture of most traits relative to Gaussian priors (de Los Campos *et al*. 2013). Sparser specifications have already been popularized in whole genome prediction (**WGP**), such as the Student *t* distribution used in BayesA (Meuwissen *et al*. 2001) and stochastic search and variable selection or SSVS (George and McCulloch 1993; Verbyla *et al*. 2009). Both specifications generally lead to far less shrinkage of large effects yet greater shrinkage of small effects compared to a Gaussian prior. In particular, the use of variable selection procedures facilitate the determination of posterior probabilities of association (**PPA**), whose control may be far more effective in maximizing both sensitivity and specificity of GWA (Fernando *et al*. 2014) compared to frequentist based inferences which require adjustments for multiple testing such as with EMMAX. Another common inferential strategy in GWA is to report the percent of variance explained by a marker or marker region (Fernando and Garrick 2013). However, point estimates of marker effects or percentage of variation explained, by themselves, do not provide formal evidence of association.

Most sparse prior WGP models have been implemented using Markov chain Monte Carlo (**MCMC**), which can be computationally expensive. Approximate analytical approaches based on the expectation–maximization (**EM**) algorithm to provide approximate maximum a posteriori (**MAP**) estimates of SNP effects have been developed to address computational limitations in these sparse prior WGP models (Meuwissen *et al*. 2009; Hayashi and Iwata 2010; Sun *et al*. 2012; Chen and Tempelman 2015). Strategies for estimating hyperparameters for MAP inference have been proposed, including those proposed by Karkkainen and Sillanpaa (2012) and Chen and Tempelman (2015), the latter adapting the average information restricted maximum likelihood (**AIREML**) algorithm for estimating hyperparameters in BayesA and SSVS specifications. These MAP implementations should also be assessed for their efficacy in GWA studies.

A pragmatic first objective in GWA is to pinpoint narrow genomic regions containing QTL rather than to specifically identify the QTL themselves, even though the latter is the ultimate goal. That is, a large number of SNP markers in a region surrounding a typically untyped QTL might be in high linkage disequilibrium (LD) with the QTL and with each other, thereby thwarting precise inference on the causal QTL. Different GWA methods may differ in the number of SNP markers inferred to have an association within a genomic region with, for example, EMMAX tending to draw associations with more SNP markers in LD with a QTL compared to use of SSVS (Guan and Stephens 2011; Goddard *et al*. 2016).

Increasingly, more GWA studies are based on inferences involving joint tests on all of the SNP markers within a narrow genomic region, recognizing that single SNP marker associations may be fraught by low statistical power or problems with multicollinearity or both (Fernando et al., 2014). Some GWA studies have been based on using several arbitrary window sizes based on either non-overlapping (Wolc *et al*. 2012; Moser *et al*. 2015; Wolc *et al*. 2016) or sliding windows (Schmid and Yang 2008). Because of the arbitrariness of fixed window sizes, whether defined by number of SNP markers or by physical length in base pairs, it is possible to split a large LD block into 2 or more separate windows, thereby making such a division seemingly suboptimal for GWA. Substantially different window lengths have been used in different studies. For example, a 5 SNP window was used for GWA based on 51,385 SNP markers in pigs (Fan *et al*. 2011), whereas a 250 kilo base (**Kb**) window was used for 287,854 SNPs from the Welcome Trust Case Control Consortium (WTCCC) human data (Wellcome Trust Case Control 2007; Moser *et al*. 2015), and a 1 mega base (**Mb**) window was used for a 24,425 SNP marker panel in chickens (Wolc *et al*. 2012). Dehman *et al*. (2015) recently proposed an approach to adaptively cluster windows of SNP markers of varying sizes based on LD relationships. That is, they performed spatially constrained hierarchical clustering of SNPs by minimizing a distance measure derived from Ward’s criterion based on LD r^2^ between SNP markers. They surmised that this procedure would estimate a suitable specification of genomic windows within each chromosome using a modified version of the gap statistic. This method has been implemented in the R package `BALD` (Dehman and Neuvial 2015).

We had three primary objectives in this study. One was to examine the potential benefits of using sparser priors (i.e., BayesA and SSVS) relative to classical (i.e., based on normality) random effects specifications and strategies for GWA under a wide range of simulated architectures. A second objective was to assess whether the choice of different fixed genomic window sizes (specifically 0.5, 1, 2, and 3 Mb), versus adaptively inferred window sizes based on LD clustering, could impact GWA performance. A final objective was to assess the relative merit of approximate MAP approaches to theoretically exact yet computationally intensive MCMC approaches based on sparse prior specifications. Our assessments are based upon SNP marker genotypes and actual and simulated phenotypes on F2 pigs deriving from a Duroc-Pietrain cross.

## METHODS and MATERIALS

### The hierarchical linear model

All analyses in this paper are based on a hierarchical linear model which be characterized by the classical mixed model specification:

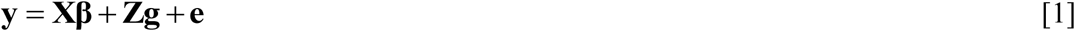

Here **y** is a *n* × 1 vector of phenotypes, **X** is a known *n* × *p* incidence matrix connecting y to the *p* × 1 vector of unknown fixed effects **β, Z** is a known *n* × *m* matrix of genotypes connecting **y** to the *m* × 1 vector of unknown random SNP marker effects **g**, and **e** is the random error vector. We also assume throughout that 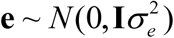 whereas 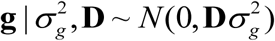 for **D** being a diagonal matrix of augmented data or variables (Chen and Tempelman 2015; Tempelman 2015). The prior specification on these diagonal elements is used to distinguish each of the competing models as described later.

For pedagogical reasons, we assume one record per individual although extensions to repeated records per individual are possible. An equivalent genomic animal effects model (VanRaden 2008) to Equation [1] can then be written as:

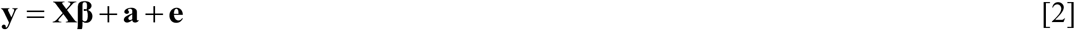
with **a = Zg** and all other terms defined previously as in [1] such that, conditionally on **D**,

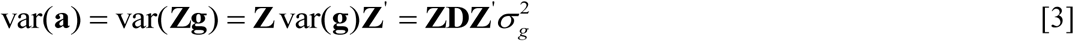

If *m* » *n*, it is generally computationally more tractable to work with the linear mixed model in Equation [2], along with the random effects specification in Equation [3], then back solve for the estimate of **g** that would be identical to those using a linear mixed model directly based on Equation [1] (Stranden and Garrick 2009).

### Models

In the simplest model, which we denote as ridge regression (RR), there is no such data augmentation (i.e. **D = I**), such that the elements of **g** are marginally distributed as independent normal (de Los Campos *et al*. 2013). Sparser distributional specifications on **g** can be constructed as mixtures of normal densities (Andrews and Mallows 1974) by simply specifying prior distributions on functions of the diagonal elements of **D**. Suppose that 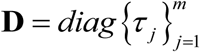 with *τ_j_* ∼ *χ™^2^*(*ν_τ_*,*ν_τ_*); then it can be demonstrated that, marginally, elements of **g** are identically and independently distributed as a scaled Student *t* with scale parameter 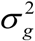 and degrees of freedom *ν_τ_* (Chen and Tempelman 2015). This model is typically referred to as BayesA (Meuwissen *et al*. 2001). Alternatively, if 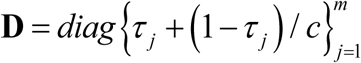 where *τ* ∼ *Bernoulli*(*π_τ_*);τ*_j_* = 0,1and *c*>>*1*, then the resulting model is Bayes SSVS in the spirit of George and McCulloch (1993). As a side-note, we use *c* = 1000 for all SSVS analyses in this paper.

As a final stage in each of the competing hierarchical models (RR, BayesA, and SSVS), we specify convenient conjugate priors wherever possible. For example, scaled inverted chi-square priors for variance components; i.e.

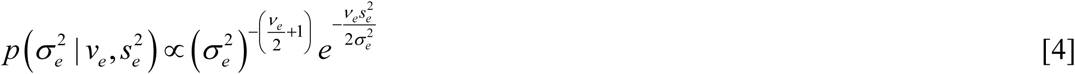
ands

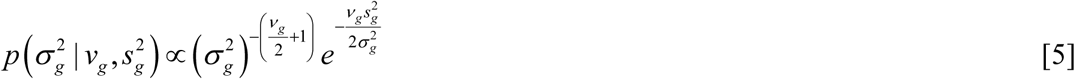
whereas we specify a Beta prior on *π_τ_* in SSVS; i.e.,

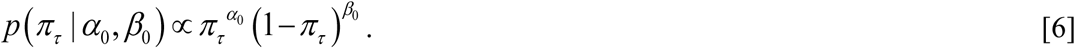

As we explain later, we arbitrarily specify *ν_τ_* as known ( *ν_τ_* =2.5), although conceptually it could also be estimated (Yang *et al*. 2015). We assume throughout that *p* (**β**)∞ 1 as *p* (**β**) is typically diffuse, although extensions to more informative specifications should be obvious. Furthermore, for all analyses in this paper, we specify Gelman’s non-informative prior (Gelman 2006) for 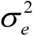 in Equation [4] based on *v_e_* = -1 and 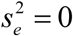 and for 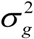 in Equation [5] based on *v_g_* = -1 and 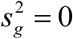. Furthermore, as per Yang and Tempelman (2012), we specify *α*_0_ = 1 and *β*_0_ = 9.

### Joint posterior density

Given the specifications above, the joint posterior density can be written as in Equation [7]

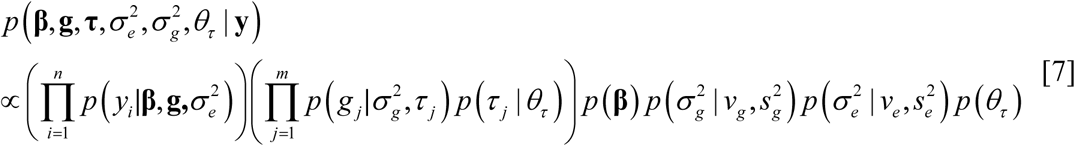
Note that *p*(*τ_j_*|*θ_τ_*) specifies the *χ*^-2^(*ν_τ_, ν_τ_*) density under BayesA (i.e., *θ_τ_* = *ν_τ_*) whereas *p*(*τ_j_*|*θ_τ_*) specifies the *Bernoulli*(*π_τ_*) density under SSVS (i.e., *θ_τ_*=*π_τ_*). Furthermore, 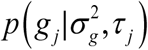 is Gaussian with null means under all three competing models but with variance 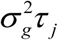 under BayesA and variance 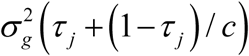 under SSVS. For RR, *τ_j_* = 1∀*j* such that 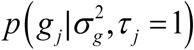 is Gaussian with common variance 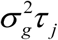.

### Algorithms

#### Markov Chain Monte Carlo

The MCMC sampling strategies that we use here for BayesA are similar to those provided in Yang and Tempelman (2012) and Yang *et al*. (2015). However, since our parameterization is slightly different, we present the full conditional densities of interest for implementing BayesA in Supplementary File S1. For similar reasons, we also provide the full conditional densities for SSVS in Supplementary File S1 as even our model differs from the model also labeled as SSVS in the genomic prediction work of Verbyla *et al*. (2009) whereas it is virtually identical to the model presented in seminal SSVS paper by George and McCulloch (1993).

#### Maximum a posterior estimation

Complete details on our MAP procedure for both BayesA and SSVS are found in Chen and Tempelman (2015). Given that our application involved *m* << *n*, we conducted MAP based inference on an equivalent animal-centric model using Equation [2] rather than based on a SNP effects model as in Equation [1]. Details on backsolving from an animal centric model to provide estimates of SNP effects are provided in Supplementary File S1. For pedagogical reasons, however, we work directly from the SNP-centric Model [1] in our subsequent developments. Conditional on **D**, the posterior variance-covariance matrix of **g**, or equivalently its prediction error variance-covariance (PEV) matrix from a frequentist viewpoint, can be written as: 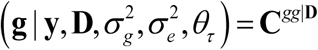. This expression can be derived from the inverse of the mixed model coefficient matrix as provided in Equation [8].

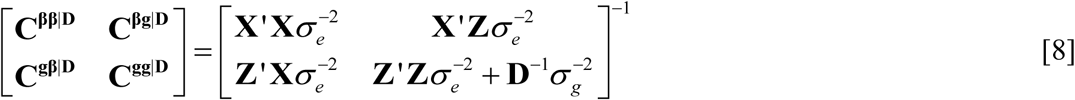

That is, **C**^gg|^**^D^** is the random by random portion of the inverse coefficient matrix in Henderson’s mixed model equations, conditional on 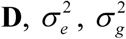 and *θ_τ_* (*θ_τ_* = *ν_τ_* for BayesA or *θ_τ_* = *π_τ_* For SSVS). As noted earlier, values for hyperparameters such as 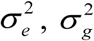 and *θ_τ_* required for Equation [8] can be determined using the REML or marginal maximum likelihood (MML) estimation strategies as described by Chen and Tempelman (2015) noting that we choose to fix *ν_τ_* in BayesA as indicated earlier.

It can be readily demonstrated (Sorensen and Gianola 2002), that asymptotically MAP (**g**) ≈ E (**g** | **y**) whereby MAP(**g**) can be iteratively determined using EM based on Newton-Raphson for maximization (M-) steps interwoven with expectation (E-) steps on elements of **D** (Chen and Tempelman, 2015). Under RR, **D** = **I** such that **C^gg^** =**C**^gg|D^ ****≈**** var **(g**|**y)** represents the posterior variance-covariance matrix of **g** conditional on 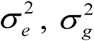 and *θ_τ_*. In fact, MAP(**g**) is synonymous with BLUP(**g**) under RR. Furthermore, **C^gg^** is synonymous with the g-component of the inverse of the negative of the Hessian of the log of the joint conditional posterior density of **β** and **g**. This posterior density is formally defined in Equation [9].

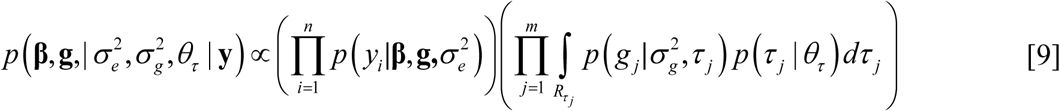

With **D** = **I**, there is no uncertainty on τ*_j_*. such the integration in Equation [9] is not necessary with **C^gg^** being directly obtainable for RR using Equation [8]. However, for BayesA and SSVS, uncertainty in **D** needs to be integrated out as per Equation [9]. An indirect strategy for asymptotically providing **C^gg^** for BayesA and SSVS is based on the strategy proposed by Louis (1982) with details provided in Supplementary File S1. We subsequently use elements of **C^gg^** to asymptotically determine key components of var (**g**|**y**) for both single SNP and window based GWA testing using MAP under all three models, noting again that MAP and BLUP are synonymous under RR.

### Conducting Genome Wide Association Analyses

#### Single SNP marker associations

We subsequently describe how we conducted GWA inference for single SNP associations based on the algorithms (MCMC vs. MAP) and models (RR, BayesA, and SSVS. With respect to inference on association on SNP *j*, EMMAX is conceptually based on subsetting out Equation [1] as follows:

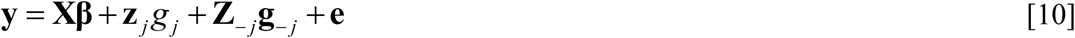

That is, **Z** is partitioned into column *j*, **z***_j_*, being the genotypes for SNP *j* and all other remaining columns in **Z***-j*. In EMMAX, *g_j_* is actually treated as fixed whereas ***g_-j_*** is treated as classically random; i.e., characterized by a Gaussian prior distribution. Writing **W**_*j*_=[**X z**_*j*_] and 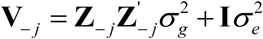, the generalized least squares (GLS) estimator *ĝ_j_* of *g_j_*, using all other markers to account for population structure, is the last element of the product 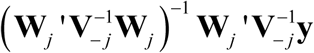. Furthermore, the corresponding standard error *se* (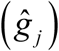) is determined by the square root of the last diagonal element of 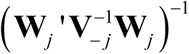. The test-statistic or “fixed effects” z-score for the EMMAX test can then be simply written as:

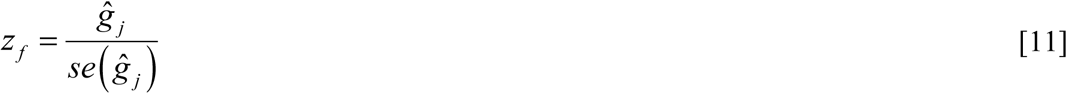
which is assumed to be *N*(0,1) under H_o_: *g_j_* = 0. The “expedited” approach (Kang *et al*. 2010) in EMMAX, that we consider in this paper, is based on approximating **V**_-_*_j_* with 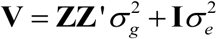 for inference of association on all SNP *j* = 1,2,…,*m;* furthermore, 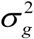 and 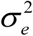 estimated only once using REML in an initial analysis that treats all SNP marker effects as random. A GWA test for a particular SNP marker *j* using EMMAX then essentially involves treating its effect jointly as both fixed and random by replacing **Z**_-_*_j_*.**g**_-_*_j_*. with **Zg** on the right side of Equation [10], implying that this double counting of *g*. as both fixed and random is trivial with large *m*.

A classical random effects test is based on treating all SNP effects, including a marker *j* of particular interest, as having a Gaussian prior such that the point estimate of the SNP substitution effect is based on fitting Equation [1] or, equivalently, backsolving from fitting Equation [3] as demonstrated by Stranden and Garrick (2009) and also in Supplementary File S1. A corresponding test statistic (*z_r_*) can be based on dividing 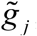, the BLUP of *g_j_* by the square root of its prediction error variance (*PEV*) where 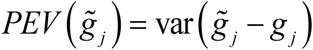 from a frequentist perspective. From a Bayesian perspective, the corresponding test statistic can be interpreted as a posterior *z*-score (Gelman *et al*. 2012) since 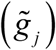 is analogous to a posterior mean (i.e., 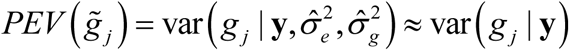) whereas the *PEV* is analogous to a posterior variance with 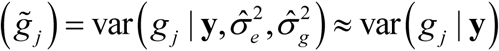. We refer to this inference strategy as **RR-BLUP**. It is important to indicate, nevertheless, that these RR-BLUP inferences are empirical Bayesian (Robinson 1991) since these posterior means and variances are typically conditioned upon REML estimates of 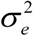 and 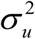. The posterior z-score (Gelman *et al*. 2012) can then equivalently derived from both frequentist and Bayesian perspectives as indicated in Equation [12].

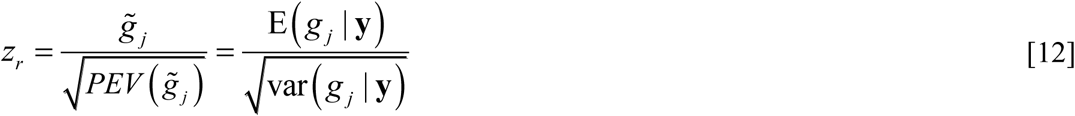

Now Gualdron Duarte *et al*. (2014), with a proof provided later by Bernal Rubio *et al*. (2016), determined that the “fixed effects” or EMMAX *z*-score, *z_f_* in Equation [11], could be equivalently derived by treating all markers as classically random, but by dividing the corresponding BLUP 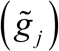 for marker *j* by the square root of its frequentist definition of variance 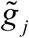 as characterized by classical mixed model theory (Searle *et al*. 1992) in Equation [13].

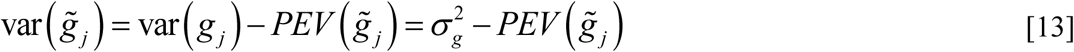

In other words, one can rewrite the fixed effects test provided in both its frequentist (numerator 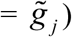) and Bayesian 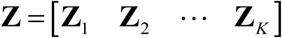) representations as in Equation [14].

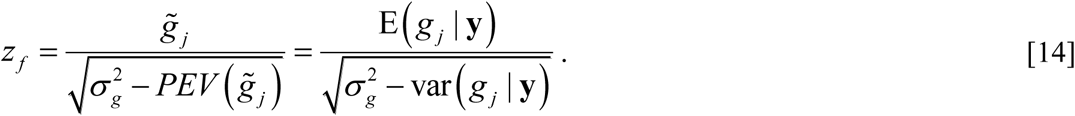

Note that the determination using Equation [14] is computationally far more tractable than that in Equation [11]. That is, Equation [14] only requires computing BLUP of **g** and its corresponding PEV in one single step determination for all *m* tests whereas Equation [11] imply *m* different mixed model analyses, each one in turn explicitly treating a different SNP marker effect as fixed.

We perceive no computationally tractable “fixed effects” test analogous to EMMAX that we could adapt for MAP based on sparser priors (BayesA and SSVS). For BayesA, for example, this would entail treating the marker of interest *j* as fixed with all other markers treated as scaled Student *t*- distributed. However, a posterior or random effects z-score test can be constructed using the MAP estimate of *g_j_* as the numerator and its asymptotic posterior standard error as the denominator, noting that MAP and the posterior mean of **g** should approach each other asymptotically. Details on deriving those asymptotic standard errors (i.e., based on deriving **C^gg^**) for use in Equation [11] for these sparse prior specifications are provided in Supplementary File S1 such that we refer to these two corresponding GWA inference strategies as MAP-BayesA and MAP-SSVS.

For SSVS based single SNP inferences using MCMC, we based inferences on the PPA for SNP marker *j* (i.e. *PPA_j_*) as in Equation [15].

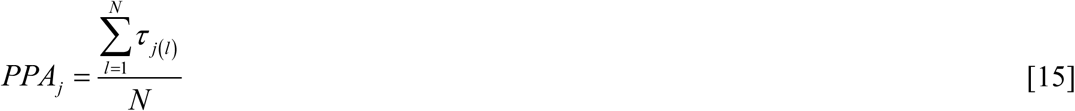

Here *N* denotes the number of MCMC cycles saved for posterior inference and *τ_j_*_(_*_l_*_)_ is a binary draw from the full conditional distribution of *τ_j_*. at MCMC cycle *l*. We denote this GWA method as MCMC-SSVS.

Since there is no variable selection inherent with BayesA under MCMC, we based single SNP inferences on a Bayesian analog to a *P*-value using

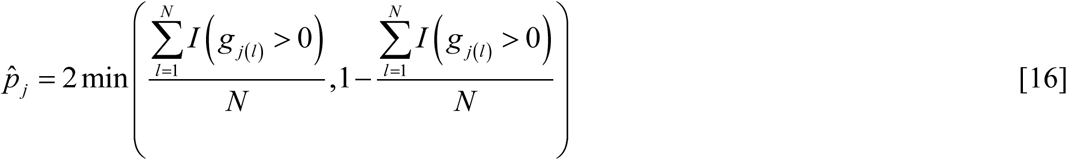

(Bello *et al*. 2010) where the indicator variable *I* (.) = 1 if the condition within the argument is true and 0 otherwise. We denote this particular GWA method as MCMC-BayesA,

#### Windows based associations

Window-based extensions to all of the above tests were also developed, some based on work previously presented above. Suppose that window *k, k* = 1,2,3,…,*K* contains *nk* markers such that **Z** can be partitioned accordingly into **Z** = [**Z**_1_ **Z**_2_… **Z***_k_*] with **Z***_k_* having *n_k_* columns, implying then that window *k* contains *n_k_* SNP markers. Similarly, the vector **g** is partitioned accordingly; i.e. 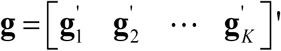 such that **g***_k_* is of dimension *nk*. × 1. Recall that we denoted 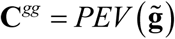. For our proposed windows-based test, the key components of **C***^gg^* can be partitioned into *K* different blocks along the block diagonal; i.e., 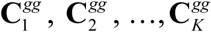 where 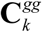 is of dimension *nk* × *nk*. The extension to a joint “fixed effects” or EMMAX like test on *n_k_* markers in window *k* involves the following extension of Equation [14].

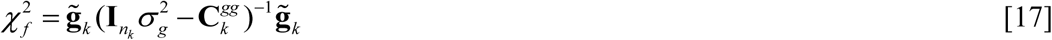

That is, it can be readily demonstrated, extending results from Bernal Rubio *et al*. (2016), that 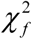 is chi-square distributed with *n_k_* degrees of freedom under H_o_: **g***_k_* = 0. The corresponding extension to a joint classical “random effects” or RRBLUP test on window *k* is provided in Equation [18]

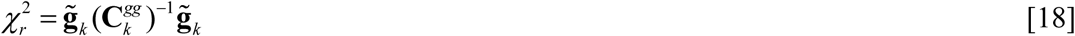
which would also be considered to be chi-square distributed with *n_k_* degrees of freedom under H_o_: **g***_k_* = 0. Similarly, one could use Equation [18] to construct the same tests for MAP-BayesA and MAP-SSVS but basing the **C^gg^** on the corresponding asymptotic posterior variance-covariance matrices as derived in Supplementary File S1.

For windows based inference using MCMC-SSVS, we simply compute the PPA for window *k* (i.e. *PPA_k_*) in Equation [19], following that presented in Fernando et al. (2014).

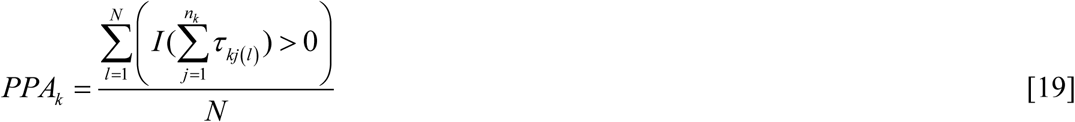

Here, τ*_kj_*_(_*_l_*_)_ defines a binary draw from the full conditional distribution of τ*_j_* for SNP marker *j* located within window *k* drawn during MCMC cycle *l*. Note then that 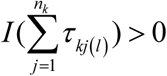 is equal to 1 when any of the draws of τ*_kj_*_(_*_l_*_)_ within window *k* are equal to 1.

For windows based GWA inference under MCMC-BayesA, we propose inferring upon the posterior probability of the proportion (*q_w_*) of the genetic variance explained by the markers in a genomic window relative to the total genetic variance as proposed by Fernando and Garrick (2013) and determined in the following manner. First note that the genotypic value that is attributed to a genomic window *k* is defined as in Equation [20].

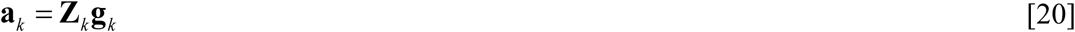

Then the variance explained by the window is defined as

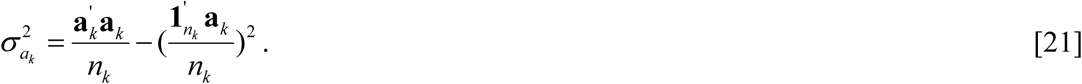

Similarly, the total genetic variance is computed as

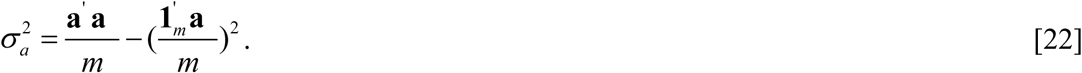

Hence, the proportion of genetic variance that is explained by marker in window *k* is defined as

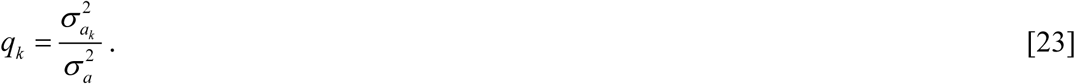

Suppose that we deem genomic windows that explain more than 1% of the total genetic variance as being of potential interest. Hence, a variable selection modification of MCMC-BayesA can be simply be based on the proportion of MCMC samples for which the genetic variance (*q_k_*) for window *k* exceeds 0.01 (Fernando and Garrick 2013). One advantage of this approach is that it can be applied to any MCMC analyses based on a model where variable selection is not explicitly specified.

### Data

#### Simulation Study

In order to compare the various models (RR, BayesA, and SSVS) and algorithms (MAP vs. MCMC), we simulated data based on the Michigan State University Pig Resource Population (MSUPRP) raised at the Michigan State University Swine Teaching and Research Farm, East Lansing, MI (Edwards *et al*. 2008). We specifically started with the SNP markers chosen for analysis by Gualdron Duarte *et al*. (2014) which included 928 Duroc-Pietrain F2 crosses. Roughly 1/3 of these pigs were directly genotyped using the Illumina Porcine SNP60 beadchip (60K) whereas the remaining F2 animals with genotyped using a lower density 9K set but whose genotypes were subsequently imputed to the 60K set (Gualdron Duarte *et al*. 2013). Edits excluded animals with more than 10% of their SNP markers missing, excluding SNP markers with more than 10% of animals missing genotypes for those markers, and excluding SNPs with minor allele frequency (MAF) below 0.01 (Gualdron Duarte *et al*. 2014). Some adjacent markers were in complete LD with each other. To circumvent multicollinearity issues, particularly its role in generating multimodality in the MCMC generated posterior densities for some SNP markers (Calus *et al*. 2015), we randomly deleted one SNP within an adjacent pair in complete LD with each other before further analyses. After invoking this edit, 43,266 SNPs remained. The original data source can be downloaded at https://msu.edu/∼steibelj/JPfiles/GBLUP.html.

To simulate different but representative genetic architectures, we generated QTL effects from three different Gamma densities with demonstrably different values of shape (γ) ranging from an effectively oligogenic density (γ = 0.18) which effectively specifies relatively much fewer QTL with large effects to an effectively polygenic Gaussian density (γ = 3.00) where most QTL have intermediate effects with symmetrically small and large effects on either side. A third intermediate value (γ = 1.48) was also chosen. A good illustration of the gamma density of QTL effects based on these three different specifications for γ is provided in Figure 1. Note that this range in γ values for QTL effects has been reported for various traits in livestock based on previous empirical work (Hayes and Goddard 2001).

**Figure 1.**
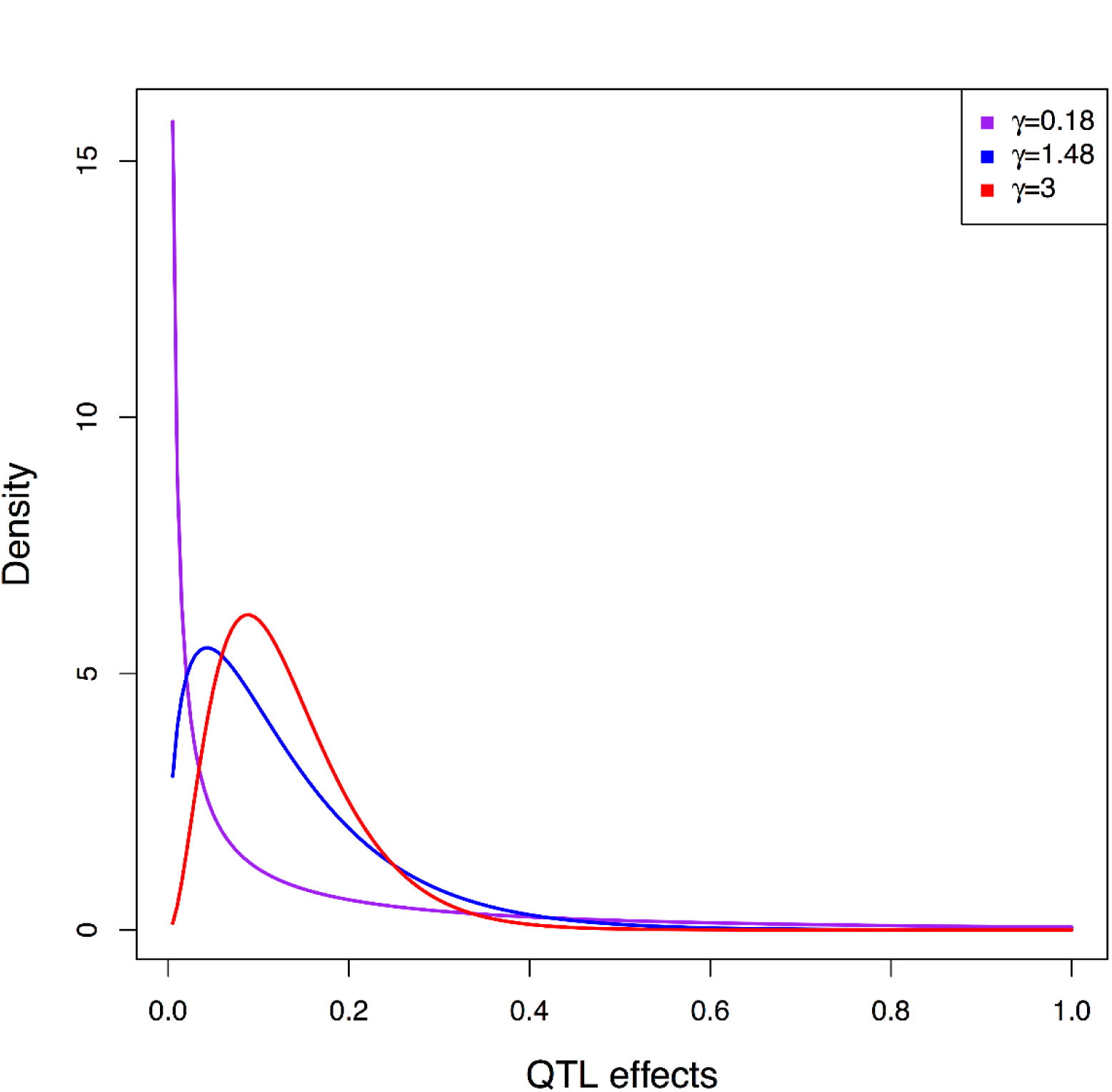
Distribution of quantitative trait loci effects under a Gamma distribution for different specifications of shape (magenta curve γ = 0.18, blue curve γ = 1.48 and red curve γ = 3.00)

In addition to the distribution of QTL effects, we also conjectured that the number of QTLs (*n_qtl_*) may also influence GWA performance such that we considered *n_qtl_* = 30, 90, or 300. Hence, we simulated 10 replicated populations under each of the 3 × 3 = 9 different scenarios pertaining to the 3 different values for each of γ and of *n_qtl_*. Each of the 90 simulated datasets were based on utilization of the 43,266 SNP marker genotypes on the *n* = 922 MSUPRP F2 pigs as previously described. Within each dataset, allelic substitution effects, **g***_qtl_*, were simulated for each of the *n_qtl_* randomly chosen SNP markers from the corresponding gamma distribution having shape γ, with a randomly chosen half of those effects multiplied by -1 as per Meuwissen *et al*. (2001). The corresponding genotypes **Z***_qtl_* for QTL on these animals were then a *n* × *n_qtl_* subset of the SNP genotype matrix **Z** such that the cumulative genetic merit or true breeding values was determined as **u***_TRUE_* = **Z***_qtl_***g***_qtl_*. Phenotypes for animals were generated based on a heritability of 0.45 as estimated for 13^th^-week tenth rib backfat from this same dataset. Only the remaining (i.e., non-QTL) marker genotypes **Z***-qtl* were used for all simulation study analyses.

In the simulation study, all parameters excluding *ν_τ_* in BayesA were estimated using both MCMC and MAP. For MCMC, we ran 200,000 iterations, discarding the first 100,000 iterations as burn-in and basing inference on saving every 10 of the remaining 100,000 cycles for a total of 10,000 samples from the posterior density. Using MAP, estimation of variance components **(θ**) for BayesA and SSVS was based on a convergence criterion of 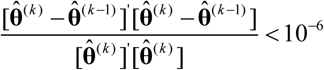. Based on our previous experience (Chen and Tempelman, 2015), we recognized that the specification of starting values in MAP-SSVS and MAP-BayesA was important for genomic prediction accuracy and, hence, likely important for GWA inferences as well. Strategies for specifying starting values for 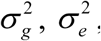, **g** and **τ** may pragmatically involve using REML and RRBLUP inferences as in Chen and Tempelman (2015) since RRBLUP is not computationally intensive. For MAP-BayesA, starting values were based on REML estimates 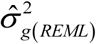 and 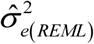 using 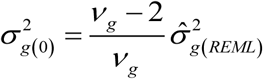 for 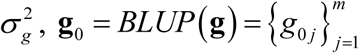 for **g** and 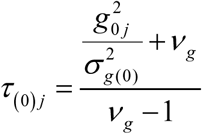 for *τ_j_, j* = 1, 2,…, *m*, based on the posterior expectation derived from its full conditional density. For MAP-SSVS, the corresponding starting values were 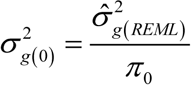 for 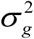 with the starting value *π*_*τ*(0)_ for *π_τ_* based, in turn, on starting values for T_;_ (i.e., SNP-specific PPA) which were determined in the following manner. First of all, EMMAX-based *P*-values for each SNP were converted to local false discovery rate (lFDR) estimates using the R package `ashr` (Stephens 2017). It has been demonstrated that these lFDR estimates, in turn, can be used to approximate PPA using PPA ≈ 1- lFDR (Stephens 2017). These approximate PPA values were then chosen as the starting values for τ*_j_* in MAP-SSVS. In turn, these starting values for τ*_j_* were used to derive the starting value for π_0_ in MAP-SSVS using the posterior expectation from its full conditional density, i.e., 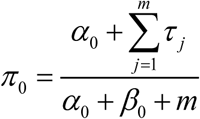. Upon convergence of variance components using the AIREML procedure outlined in Chen and Tempelman (2015), convergence of MAP-based solutions to **g** were based on the same criteria.

Single SNP marker inferences were based on the procedures outlined previously; i.e. for MAP by comparing *z_r_* in Equation [12] for the random effect tests for RRBLUP, MAP-BayesA, and MAP-SSVS and *z_f_* for the EMMAX test in Equation [11] to a standard normal distribution. No adjustment for multiple testing were invoked for the shrinkage based procedures (RRBLUP, MAP-BayesA, and MAP-SSVS) as per Gelman *et al*. (2012) whereas a Bonferonni adjustment based on the number of markers was invoked for EMMAX. Furthermore, the posterior means provided in Equations [15] and [16] were used for GWA under MCMC-SSVS and MCMC-BayesA, respectively. Since the remaining genotypes **Z***-qtl* did not include the simulated QTL, SNP markers were declared as true positives if the QTL was located between it and its closest SNP neighbor on either side.

Window based inference was based on the procedures outlined previously; i.e. for MAP by computing 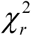 in Equation [18] for the random effect tests using RRBLUP, MAP-BayesA, and MAP-SSVS and 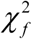 for fixed effects test in Equation [17] under EMMAX. These test statistics were compared to a chi-square distribution with degrees of freedom *n_k_*. Similar to single marker tests, no multiple testing adjustments were incurred for 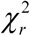 whereas a Bonferonni adjustment based on the total number of genomic windows was invoked for 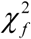. Furthermore, GWA was based on the PPA that *q_k_* > 0.01 as provided in Equation [23] for MCMC-BayesA and on the PPA for MCMC-SSVS as provided in Equation [19].

For windows-based inference, four alternative fixed window sizes were chosen: 0.5, 1, 2, or 3 Mb. The genome map used was the *Sus Scrofa* build 10.2 (http://www.ensembl.org/Sus_scrofa/Info/Index). Also, as per Moser *et al*. (2015), two different within-chromosome starting positions (starting at location 0 or 0.25 Mb for window size 0.5; starting at 0 or 0.5 Mb location for window sizes 1 Mb; starting at 0 or 1 Mb location for window sizes 2Mb; and starting at 0 or 1.5 Mb location for window sizes 3Mb) for each chromosome were chosen to partly counteract the chance effect of different LD patterns being associated with non-overlapping windows. Finally, adaptive window sizes based on clustering SNP by LD *r^2^*were also determined implementing the `BALD` R package (Dehman and Neuvial 2015) using the procedure described by Dehman *et al*. (2015).

The relative performance of all methods and models were based on receiver operating characteristic (ROC) curves. In a ROC curve, the true positive rate (TPR) is plotted against the false positive rate (FPR) for each competing method (Metz 1978). We were more specifically interested in the partial area under the curve up until a FPR= of 5% (pAUC05) so as to not include somewhat irrelevant ROC regions with low levels of specificity (Ma *et al*. 2013). A perfect classifier would have a pAUC05 of 0.05 × 1 = 0.05 whereas a random classifier would have a pAUC05 of 0.05^2^/2= 0.00125. We subsequently rescaled all pAUC05 measures by 0.00125^-1^ such that a random classifier is rescaled to a relative pAUC05 = 1. We used the R package ROCR (Sing *et al*. 2005) to obtain replicate-specific ROC curves and pAUC05 for each of the 10 replicated datasets for each method and window specification within each *n_qtl_* and γ combination. For each window specification, specific comparisons between methods were based on using the logarithm of pAUC05 as the response variable in a mixed model ANOVA with methods, *n_qtl_* and γ and all of their interactions included as fixed effects and population replicate (nested within *n_qtl_* and γ) as a random effect blocking factor. For windows-based inferences based on fixed window sizes, replicate-specific pAUC05 values were averaged over the two different starting positions as previously noted. Mean log(pAUC05) estimates were backtranformed (i.e. anti-logged) to the original scale for reporting. Overall marginal means were separated using Tukey’s test whereas comparisons between methods were sliced out using ANOVA t-tests for each value of *n_qtl_* or γ if the corresponding interaction between these factors with methods were significant (*P*<0.05). We are also conjecture that window size might actually influence of the power of detecting QTL using the same method; therefore, we conducted separate tests comparing pAUC05 for each of the different window sizes, including adaptively chosen windows based on `BALD`, separately within each method.

#### MSUPRP data

We also compared all models and algorithms on 13-week tenth rib backfat (mm) within the MSUPRP data as per Gualdron Duarte *et al*. (2014). Sex, contemporary group, and age of slaughter were treated as fixed effects (i.e., **β**). We compared each of the six competing methods, computing either PPA or *P*-values in the same manner as in the simulation study. For MCMC-BayesA and MCMC-SSVS, we ran a total of 1 million MCMC iterations based on 500,000 burn in iterations and 500,000 iterations post burn-in saving every 10 iterations such that posterior inference was based on 50,000 random draws from the posterior distribution. Since we did not know the true positions of the causal QTL for this trait, GWA inferences were compared between the various methods, based on PPA for MCMC-BayesA and MCMC-SSVS, *P*-values for RRBLUP, MAP-BayesA, and MAP-SSVS, and Bonferroni adjusted P-values for EMMAX.

## RESULTS

### Simulation Study

Overall mean comparisons between methods for pAUC05 based on single SNP inferences are provided in Table 1, noting that two-way interactions were not detected (*P* > 0.05) between methods with γ or with *n_qtl_*. There was no evidence of a sizeable difference between any of the methods given that pAUC05 ranged from 2.52 to 2.77 times that for a random classifier, although MCMC-BayesA ranked lowest.

**Table 1:**
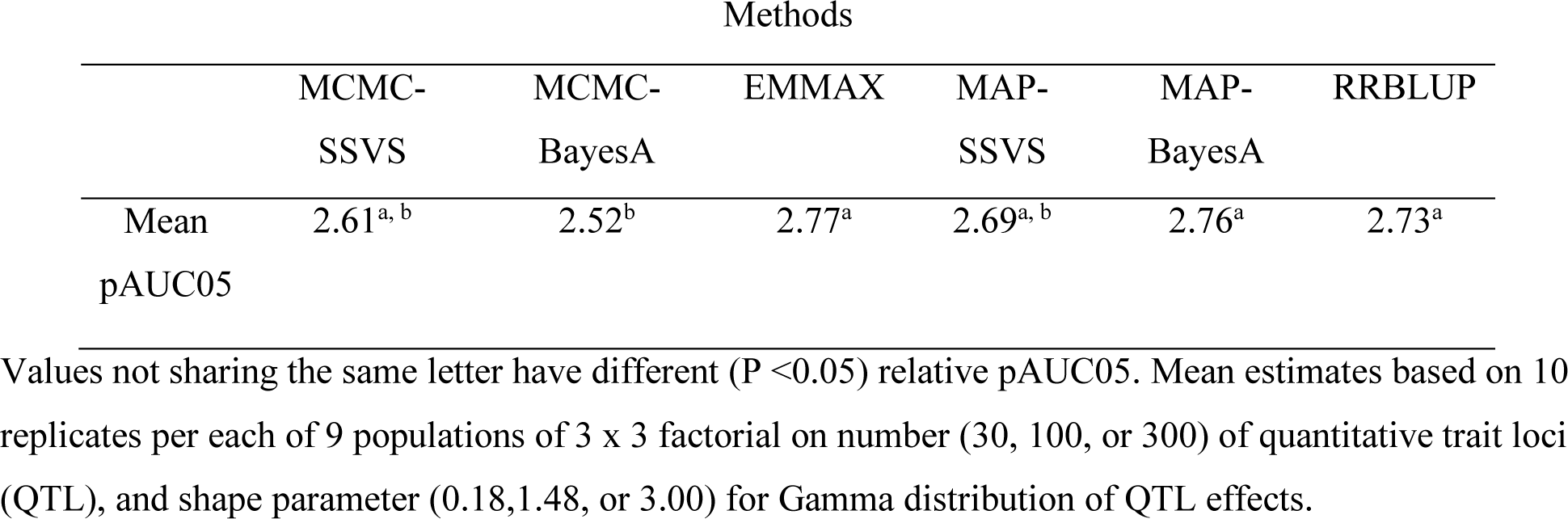
Overall mean relative (random classifier = 1) partial areas under a receiving operating characteristic curve up until a false positive rate of 5% (pAUC05) for different methods on single SNP associations

|  | Methods |  |  |  |  |  |
| --- | --- | --- | --- | --- | --- | --- |
|  | MCMC-<br>SSVS | MCMC-BayesA | EMMAX | MAP-SSVS | MAP-BayesA | RRBLUP |
| Mean<br>pAUC05 | 2.61<sup>a, b</sup> | 2.52<sup>b</sup> | 2.77<sup>a</sup> | 2.69<sup>a, b</sup> | 2.76<sup>a</sup> | 2.73<sup>a</sup> |

Values not sharing the same letter have different ( $P < 0.05$ ) relative pAUC05. Mean estimates based on 10 replicates per each of 9 populations of 3 x 3 factorial on number (30, 100, or 300) of quantitative trait loci (QTL), and shape parameter (0.18, 1.48, or 3.00) for Gamma distribution of QTL effects.

For fixed 1Mb window sizes (Table 2), the two-way interactions between method and γ and between method and *n_qtl_* were both significant (*P* < 0.0001). Therefore, methods were compared separately for each different value of γ and of *n_qtl_*. Nevertheless, MCMC-SSVS and MCMC-BayesA had the largest pAUC05 (*P* < 0.05) for each different value of γ and of *n_qtl_* as well as overall. EMMAX generally followed MCMC-SSVS and MCMC-BayesA with MAP-SSVS, MAP-BayesA and RRBLUP being the worst performing methods. Most notably, these latter three methods generally did worse than a random classifier (i.e. pAUC05 < 1) except for MAP-SSVS at *n_qtl_* = 30.

**Table 2:** Least squares mean relative (random classifier = 1) partial areas under a receiving operating characteristic curve up until a false positive rate of 5% (pAUC05) for different methods for associations based on genomic windows of length 1Mb. Comparisons are made within different specifications of shape parameter (γ) for Gamma distribution of quantitative trait loci (QTL) and number of QTL (*nqti*)

| Factors | Methods |  |  |  |  |  |
| --- | --- | --- | --- | --- | --- | --- |
|  | MCMC-SSVS | MCMC-BayesA | EMMAX | MAP-SSVS | MAP-BayesA | RRBLUP |
| Shape $\gamma$ | | | | | | |
| 0.18 | 2.82<sup>a</sup> | 2.75<sup>a</sup> | 1.78<sup>b</sup> | 0.74<sup>c, *</sup> | 0.63<sup>c, *</sup> | 0.48<sup>d, *</sup> |
| 1.48 | 4.22<sup>a</sup> | 4.16<sup>a</sup> | 2.54<sup>b</sup> | 0.69<sup>c, *</sup> | 0.38<sup>d, *</sup> | 0.28<sup>e, *</sup> |
| 3 | 4.63<sup>a</sup> | 5.01<sup>a</sup> | 2.47<sup>b</sup> | 0.67<sup>c, *</sup> | 0.40<sup>d, *</sup> | 0.24<sup>e, *</sup> |
| $n_{qtl}$ | | | | | | |
| 30 | 6.81<sup>a</sup> | 7.28<sup>a</sup> | 3.63<sup>b</sup> | 1.69<sup>c</sup> | 0.67<sup>d, *</sup> | 0.31<sup>e, *</sup> |
| 90 | 3.89<sup>a</sup> | 3.78<sup>a</sup> | 2.14<sup>b</sup> | 0.61<sup>c, *</sup> | 0.47<sup>c, d, *</sup> | 0.37<sup>d, *</sup> |
| 300 | 2.08<sup>a</sup> | 2.08<sup>a</sup> | 1.42<sup>b</sup> | 0.33<sup>c, *</sup> | 0.31<sup>c, *</sup> | 0.29<sup>c, *</sup> |
| Overall | 3.81<sup>a</sup> | 3.86<sup>a</sup> | 2.23<sup>b</sup> | 0.70<sup>c, *</sup> | 0.46<sup>d, *</sup> | 0.32<sup>e, *</sup> |

Values not sharing the same letter within a row have different ( $P < 0.05$ ) relative pAUC05 within the row. \* indicates the corresponding method is not better than a random classifier. Mean estimates based on 10 replicates per each of 9 populations of 3 x 3 factorial on number (30, 100, or 300) of quantitative trait loci (QTL), and shape parameter (0.18, 1.48, or 3.00) for Gamma distribution of QTL effects.

Table 3 highlights the comparisons between the various methods using the adaptive window sizes inferred by `BALD`. Here, the two-way interaction between method and *n_qtl_* was important (*P* < 0.05) whereas the two-way interaction between method and γ was not; hence, we just compared different methods within each different value of *n_qtl_*. As with the 1Mb window inferences, MCMC-SSVS and MCMC-BayesA had the highest pAUC05, followed by EMMAX within each different value of *n_qtl_* such that these same rankings were found overall as well. Again, we found that MAP-SSVS, MAP-BayesA, and RRBLUP had lower pAUC05 compared to a random classifier except for MAP-SSVS when *n_qtl_* = 30.

**Table 3:** Least squares mean relative (random classifier = 1) partial areas under a receiving operating characteristic curve up until a false positive rate of 5% (pAUC05) for different methods for associations based on genomic windows adaptively chosen by the `BALD` software package. Comparisons are made within different specifications of number of quantitative trait loci (*nqti*)

| $n_{qtl}$ | Methods | | | | | |
| --- | --- | --- | --- | --- | --- | --- |
|  | MCMC-SSVS | MCMC-BayesA | EMMAX | MAP-SSVS | MAP-BayesA | RRBLUP |
| 30 | 8.83<sup>a</sup> | 9.03<sup>a</sup> | 3.57<sup>b</sup> | 1.76<sup>c</sup> | 0.87<sup>d,*</sup> | 0.29<sup>e,*</sup> |
| 90 | 5.34<sup>a</sup> | 4.98<sup>a</sup> | 2.13<sup>b</sup> | 0.8<sup>c,*</sup> | 0.50<sup>d,*</sup> | 0.38<sup>d,*</sup> |
| 300 | 3.89<sup>a</sup> | 3.17<sup>a</sup> | 1.30<sup>b</sup> | 0.66<sup>c,*</sup> | 0.62<sup>c,*</sup> | 0.58<sup>c,*</sup> |
| Overall | 5.68<sup>a</sup> | 5.22<sup>a</sup> | 2.15<sup>b</sup> | 0.97<sup>c,*</sup> | 0.65<sup>d,*</sup> | 0.40<sup>e,*</sup> |

Values not sharing the same letter within a row have different ( $P < 0.05$ ) relative pAUC05 within the row. \* indicates the corresponding method is not better than a random classifier. Mean estimates based on 10 replicates per each of 9 populations of 3 x 3 factorial on number (30, 100, or 300) of quantitative trait loci (QTL), and shape parameter (0.18, 1.48, or 3.00) for Gamma distribution of QTL effects.

We were also interested in pAUC05 comparisons between different window length specifications. Recognizing that the interaction between method and window length was important in our joint analysis involving all simulated datasets, we choose to focus on window length comparisons separately within each of MCMC-SSVS, MCMC-BayesA, and EMMAX (Table 4), given that all other methods performed worse than random classifier with windows based inference. For EMMAX, single SNP inferences has significantly larger pAUC05 compared to inferences based on the longer genomic windows (2 and 3 Mb) with inference based on adaptively determined windows using `BALD` and shorter genomic windows (0.5Mb and 1Mb) being intermediate in their performance. Conversely, for both MCMC-BayesA and MCMC-SSVS, single SNP inference had the lowest pAUC05 whereas adaptively determined window selection based on `BALD` yielded the highest pAUC05 with fixed window inferences being intermediate in their performance. In fact, the best overall performance was based on using the two MCMC based methods with adaptively determined windows with a pAUC05 being over 5 times greater than that of a random classifier.

**Table 4:** Least squares mean relative (random classifier = 1) partial areas under a receiving operating characteristic curve up until a false positive rate of 5% (pAUC05) between different window sizes within each of EMMAX, MCMC-BayesA, and MCMC-SSVS.

| EMMAX |  | MCMC-BayesA |  | MCMC-SSVS |  |
| --- | --- | --- | --- | --- | --- |
| Window | pAUC05 | Window | pAUC05 | Window | pAUC05 |
| Single SNP | 2.76<sup>a</sup> | Single SNP | 2.52<sup>c</sup> | Single SNP | 2.61<sup>c</sup> |
| 0.5Mb | 2.43<sup>a, b</sup> | 0.5Mb | 3.77<sup>b</sup> | 0.5Mb | 3.65<sup>b</sup> |
| 1Mb | 2.23<sup>b, c</sup> | 1Mb | 3.86<sup>b</sup> | 1Mb | 3.81<sup>b</sup> |
| 2Mb | 1.95<sup>c</sup> | 2Mb | 3.93<sup>b</sup> | 2Mb | 3.94<sup>b</sup> |
| 3Mb | 1.85<sup>c</sup> | 3Mb | 3.93<sup>b</sup> | 3Mb | 4.04<sup>b</sup> |
| Adaptive | 2.15<sup>b, c</sup> | Adaptive | 5.22<sup>a</sup> | Adaptive | 5.67<sup>a</sup> |

Values not sharing the same letter within a column have different ( $P < 0.05$ ) relative pAUC05 within the column. \* indicates the corresponding method is not better than a random classifier. Mean estimates based on 10 replicates per each of 9 populations of 3 x 3 x 5 factorials on number (30, 100, or 300) of quantitative trait loci (QTL), shape parameter (0.18, 1.48, or 3.00) for Gamma distribution of QTL effects, and genomic region size (single SNP, 0.5Mb, 1Mb, 2Mb, 3Mb or adaptively determined) for genome wide association.

### MSUPRP Data

Manhattan plots based on single SNP associations for 13-week tenth rib backfat (mm) in MSUPRP are provided in Figure 2. The statistically most significant marker identified by EMMAX was SNP label ALGA0104402 (*P* = 2.36e-10) at location 136.0844Mb in Chromosome 6, marking the same location identified as being most significantly associated with this trait by Gualdron Duarte *et al*. (2014). Another 11 nearby statistically significant markers ranged in location from 132.60Mb to 138.24Mb with 1 marker (SNP label MARC0035827) at 122.36Mb on Chromosome 6 being also statistically significant using EMMAX. For MCMC-SSVS, the marker (SNP label ALGA0122657) located at 136.0786Mb on Chromosome 6 had the highest PPA of 0.487 and was adjacent to the most significant marker ALGA0104402 as identified by EMMAX. MCMC-SSVS also inferred its second largest PPA=0.227 with SNP marker ALGA0104402. Hence, the top 2 SNP markers identified by MCMC-SSVS and EMMAX were the same, albeit their order of importance was reversed. Although the most significant single SNP associations were also determined within this same region for each of the four other methods, their levels of significance were clearly not important except perhaps for MAP-SSVS which started to approach statistical significance with SNP label MARC0035827 (P=0.08).

**Figure 2.**
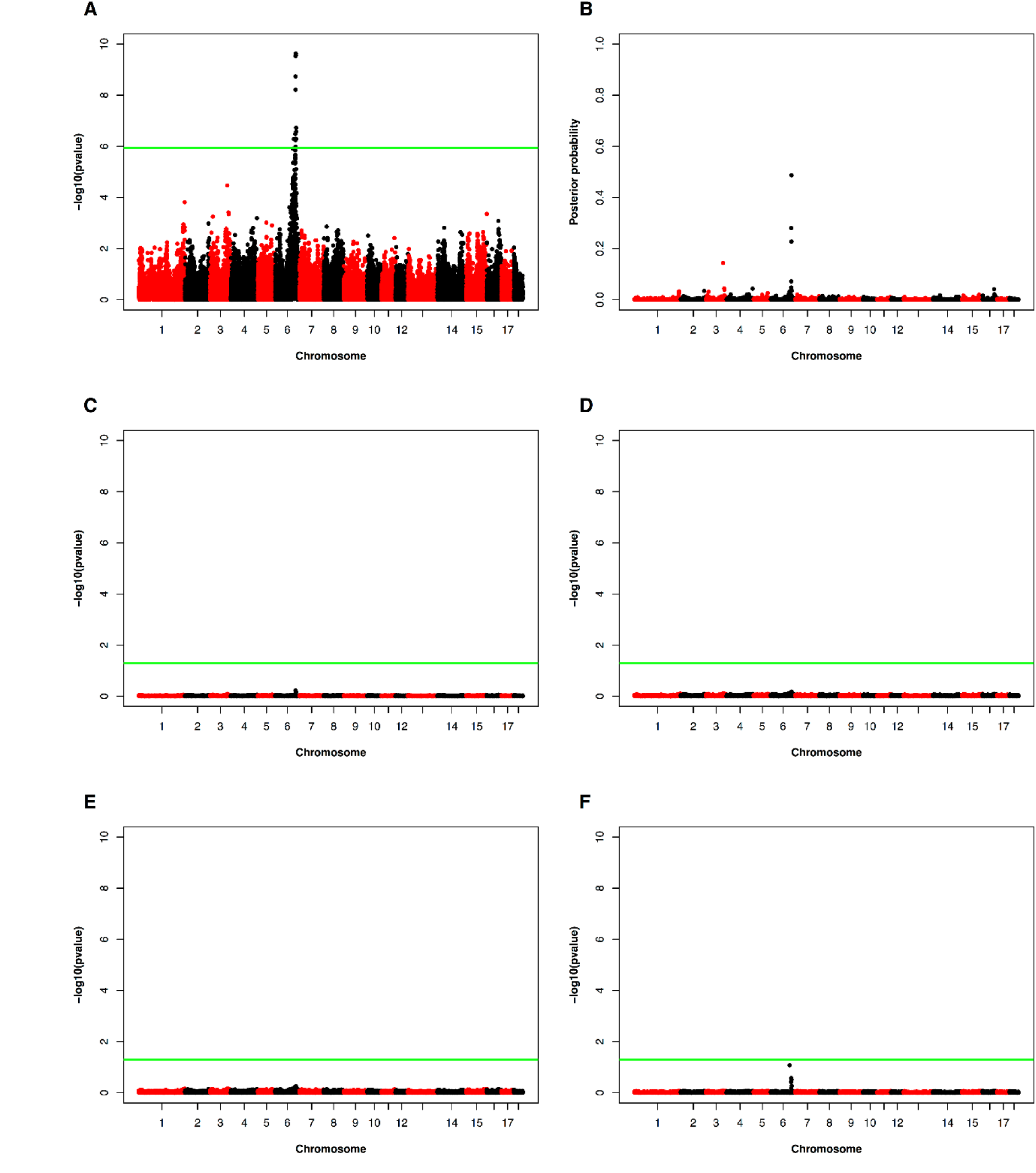
Manhattan plots for single SNP analysis on 13^th^ week 10^th^ rib backfat in Duroc Pietrain F2 cross (*n* = 922 pigs) based on different methods (Panel A: EMMAX, Panel B: MCMC-SSVS, Panel C: MCMC-BayesA, Panel D: RRBLUP, Panel E: MAP-BayesA and Panel F: MAP-SSVS)

For windows-based inference, we focused on the adaptively chosen window strategy based on LD using `BALD` (Figure 3). For EMMAX, the most significant window (P = 9.36e-08) ranged from 134.17Mb to 134.75Mb on Chromosome 6. Although this region did not contain any markers that were statistically significant based on single SNP based inferences, it was very close to a marker (SNP label ASGA0029653) at 134.14Mb that was deemed to be statistically significant in Figure 2. Four other windows on Chromosome 6 were also significant, covering regions 129.70-131.35Mb, 132.87-134.14Mb, 135.19-136.84Mb and 136.97-137.32Mb. These windows included some statistically significant or nearly significant markers based on single SNP inferences in Figure 2. Using MCMC-SSVS, the most significant window (Window 909) covered 135.19-136.84Mb with a PPA = 0.722; this window also contained the most significant markers based on single SNP inferences using EMMAX and MCMC-SSVS in Figure 2. Window 905 had the second highest PPA = 0.477 and ranged in location from 132.87-134.14Mb with all other windows having smaller PPA (< 0.2). A LD heatmap of the genomic region containing both windows are provided in Figure 4, indicating that some SNP markers in Window 905 are in relatively high LD with markers in Window 909. These two windows also had the highest PPA under MCMC-BayesA being 0.459 and 0.553 respectively. For RRBLUP, MAP-BayesA and MAP-SSVS, no window was deemed to be statistically significant (*P*>0.05).

**Figure 3.**
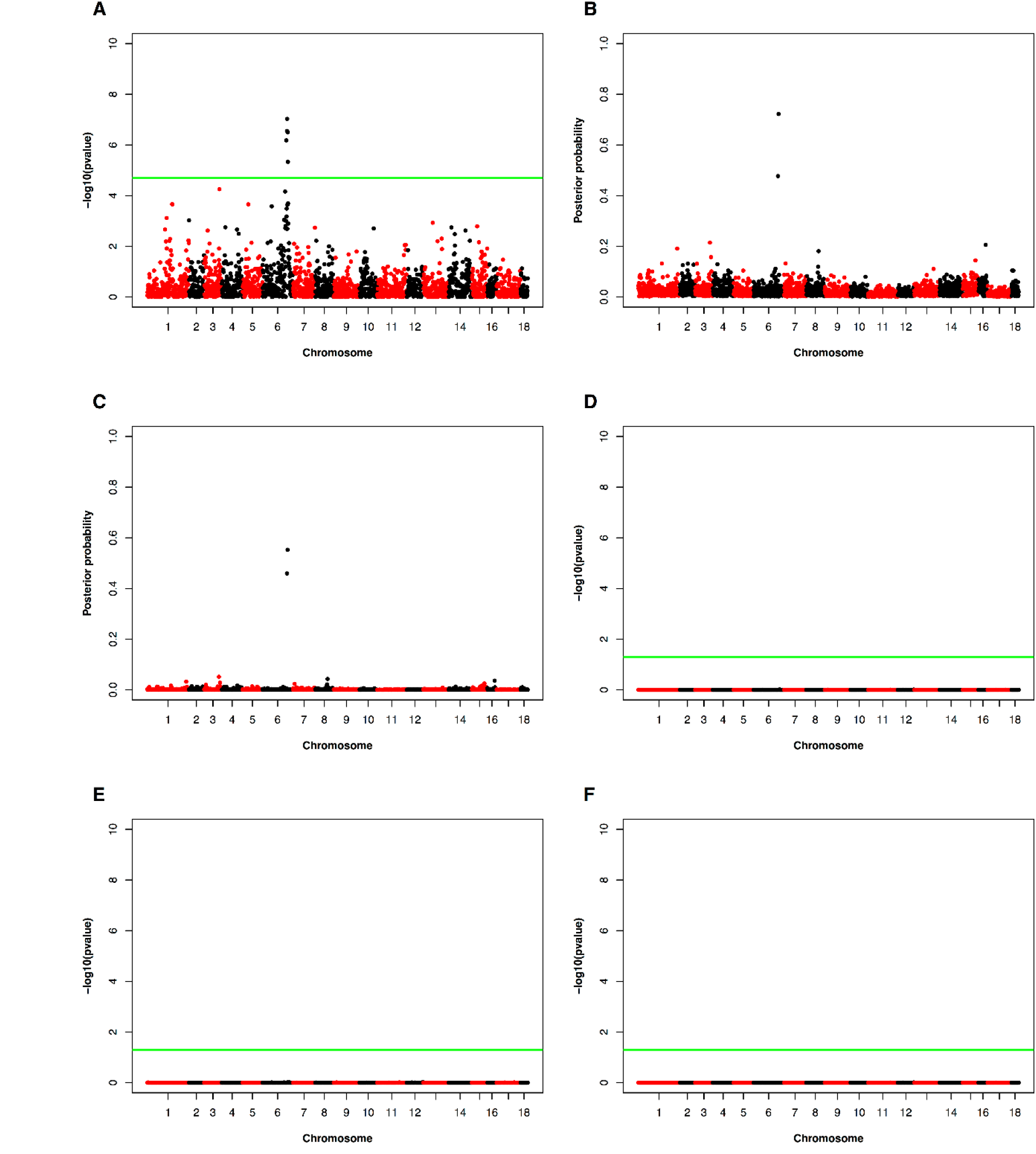
Manhattan plots for genomic window based associations on 13^th^ week 10^th^ rib backfat in Duroc Pietrain F2 cross (n = 922 pigs) based on different methods (Panel A: EMMAX, Panel B: MCMC-SSVS, Panel C: MCMC-BayesA, Panel D: RRBLUP, Panel E: MAP-BayesA and Panel F: MAP-SSVS) under adaptive window inference.

**Figure 4.**
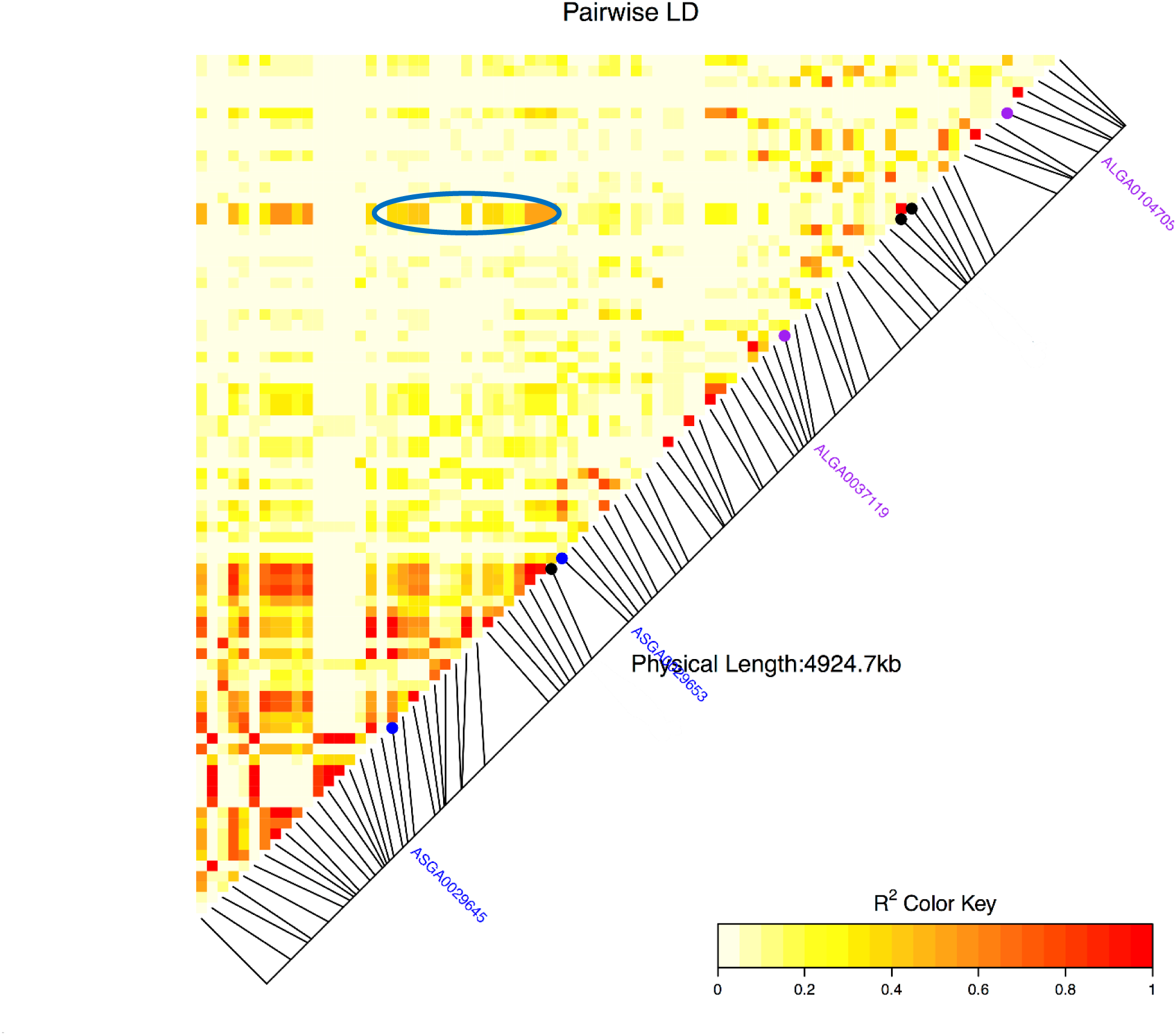
Linkage disequilibrium (*r*^2^ metric) heatmap for genomic region containing Windows 905 - 909 on Chromosome 6 as adaptively determined by `BALD` software. Blue dots are starting and ending points for window 905 whereas purple dots are starting and ending points for window 909. Black dots are the 3 markers at 133.9292Mb, 136.0786Mb and 136.0844Mb that are top 3 SNPs by MCMC-SSVS. The blue oval is used to highlight a pocket of higher *r*^2^ measures SNP markers in windows 905 and 909.

## DISCUSSION

The objectives of our study were multifaceted in that we wished to very broadly address the impact of a) prior specifications on marker effects, b) single marker associations versus associations based on different specifications for genomic windows and c) of computationally tractable but analytical approximations for GWA inference based on sparse priors. Although our simulation study was based on genotypes derived from a specific population (MSUPRP), a wide variety of potential genetic architectures were constructed on top of that framework based on different degrees of skewness of a Gamma distribution via different specifications of the shape parameter (γ) for QTL effects as well as the number (*n_qtl_*) of QTL.

A majority of GWA studies have been conducted using single SNP inferences (Goddard and Hayes 2009; Visscher *et al*. 2012; Goddard *et al*. 2016). In this specific context, we determined that the difference in pAUC05 between all methods were relatively small and unimportant even though MCMC-BayesA had significantly lower pAUC05 and hence worse GWA performance. However, for all windows based analyses, MCMC-BayesA and MCMC-SSVS had significantly greater pAUC05 than all other methods across all combinations of γ and *n_qtl_*, regardless of window size and whether these window sizes were fixed or adaptively inferred based on LD using the `BALD` software package. Conceptually, MCMC-BayesA might have even outperformed MCMC-SSVS for windows-based GWA as our comparisons may have been influenced by the arbitrariness of using 1% as a threshold for percentage of total genetic variance explained by a window when determining the PPA under MCMC-BayesA. That is, proper specification of such a threshold is likely to be density dependent. Admittedly, a BayesB like implementation (Meuwissen *et al*. 2001) could have captured the best features (i.e. variable selection and heavy-tailed priors) of both BayesA and SSVS. EMMAX typically ranked third whereas MAP implementations of BayesA and SSVS as well as RRBLUP did much more poorly for windows based association. The latter was not too surprising since previously Gualdron Duarte *et al*. (2014) also determined that RRBLUP was extremely conservative for GWA in this same dataset. Furthermore, this liability of RRBLUP has been noted by others including Hayes (2013). We noted that the median and mean lengths for windows adaptively chosen by `BALD` software were 0.59Mb and 0.91Mb (Panel A in Figure S1 in Supplementary File S2), respectively, such that it was reasonable to expect adaptively chosen windows to lead to an GWA performance closest to inferences based on either based on the 0.5Mb or 1Mb fixed window sizes as we did observe for the two MCMC based procedures.

What was initially surprising to us was that the pAUC05 for the analytical “shrinkage”- based procedures, namely RRBLUP, MAP-SSVS and MAP-BayesA, under windows based inference was often worse than that of a random classifier (i.e. pAUC05<1). This, at first, seemed counterintuitive to us. Hence, we briefly investigated a scenario where the number of SNP markers per window was fixed to be 10 rather than basing window sizes on a fixed physical distance. Basing genomic windows on a fixed number of SNPs has been a strategy also considered elsewhere (Zhang *et al*. 2016). In our particular case, the average length of a 10 SNP window was 0.51 Mb such that one might anticipate that inference based on 10 SNP marker windows might be comparable to using inference based on fixed 0.5 Mb length windows. Nevertheless, we determined that 10 SNP windows based inference lead to a ROC performance that was at least better than a random classifier for each of RRBLUP, MAP-SSVS and MAP-BayesA (Figure S2 in Supplementary File S2), conversely to what we observed previously to windows based on any fixed physical distance. This contrast in pAUC05 performance between fixed physical distance and fixed number of markers could be explained as follows. For the vast majority of windows based on either scenario (fixed number of markers or fixed physical distance), the P-values for the chi-square tests of these shrinkage based procedures were very large (i.e., *P* > 0.85). With inference based on a fixed number of SNP markers per window and random assignment of QTL to these markers, it was reasonable to expect that the pAUC05 of any of these procedures should be at least as large as a random classifier. However, with inference based on fixed physical distance in Mb or even adaptively determined based on LD relationships, the number of SNP markers and hence the degrees of freedom for each window-specific chi-square test is highly variable, ranging from 1 to 35 with 0.5Mb windows, for example. Hence regions with few markers are more likely to have smaller *P*-values than regions with many markers by nature of a greater penalty incurred with a larger degrees of freedom chi-square test statistic. Furthermore, lower P-value regions with fewer markers are also less likely to contain a QTL because of random assignment of QTL to markers throughout the genome such that regions with the smallest *P*-values would more likely include a greater than expected number of false positive results relative to a random classifier.

One possible strategy to mitigate this problem is through use of a likelihood ratio test for the variance component characterizing the variance attributable to markers within a window can be considered for EMMAX or the MAP based approaches as then the degrees of freedom for that test does not depend on the number of markers in that window (Wu *et al*. 2010; Wang *et al*. 2013). Gualdron Duarte *et al*. (2014) present details for such a likelihood ratio test; nevertheless, this approach requires one to refit the entire model each time that a particular window is being tested and hence can be computationally challenging.

We specifically determined that adaptive window specifications based on `BALD` worked best for both MCMC-BayesA and MCMC-SSVS with significantly higher mean pAUC05 than inferences based on fixed window lengths or single SNP markers. In fact, there was no evidence of differences in pAUC05 between GWA associations based on windows of constant sizes ranging from 0.5 to 3Mb when using either MCMC-BayesA or MCMC-SSVS. Hence adaptive window clustering based on LD measures seems to be an important factor to consider when partitioning genomic windows, at least for Bayesian sparse prior specifications.

We have previously established that starting values are important for MAP-SSVS and MAP-BayesA (Chen and Tempelman 2015); in fact, we then demonstrated that starting marker effects at null values was very suboptimal, even though that is a common strategy for genomic prediction methods based on the use of the EM algorithm (Meuwissen *et al*. 2009; Karkkainen and Sillanpaa 2012). As we adapted in this study, a practical strategy is to base starting values on RRBLUP and genomic REML as we conducted in this study although we worried as to how suboptimal that might be, recognizing MAP estimates are asymptotic i.e., MAP(**g**) → E(**g** | **y**) only as *n* and such that *n* >> *m*. Being components of the joint modal estimator of **β** and **g**, MAP(**g**) is particularly susceptible to the specification of starting values when *m* >> *n*, as it becomes then increasingly likely that the joint posterior density of **β** and **g** is multimodal. In some classical animal breeding models used for non-Gaussian responses, asymptotic MAP estimators work well, i.e., MAP (**g**) → E (**g**|**y**), when *n* » *m* (Kizilkaya *et al*. 2002) where *m*, more generally, refers to the number of random effects. Conversely, MAP may be expected to perform very poorly when *m* » *n* (Ramirez-Valverde *et al*. 2001). In the context of GWA, this asymptotic inference problem of MAP would only be expected to further exacerbate with much denser SNP marker panels (i.e., increasing m) as used in many GWA studies.

To further assess whether starting values based on RRBLUP and genomic REML estimates might lead to suboptimal GWA inferences, we also based starting values for MAP-SSVS and MAP-BayesA on posterior mean estimates derived from their MCMC counterparts, focusing only, however, on single SNP and adaptive window inference. We recognize that this would not be a practical MAP strategy as once MCMC based inferences are obtained, then asymptotic MAP based inferences would not have any extra value. As anticipated from our previous genomic prediction work (Chen and Tempelman 2015), using MCMC based starting values for MAP-SSVS lead to a larger pAUC05 compared to the use of RRBLUP or genomic REML starting values except for no evidence of a difference at *n_qtl_* = 300 (Table S5 in Supplementary File S2). However, for adaptively determined windows, even MAP-SSVS inferences based on MCMC based starting values were no better than a random classifier except for when *n_qtl_* =30. Similar results for comparing different sets of starting values (MCMC-BayesA vs BLUP) for MAP-BayesA are provided in Table S6 in Supplementary File S2. These supplementary results further illustrate how precarious is the use of MAP based procedures for Bayesian regression GWA analyses; again, we would believe the sensitivity of MAP to starting values would only be greater with the use of high density marker panels.

As our GWA inference for MCMC-SSVS was based on PPA (i.e. Prob(τ*_j_* = 1|**y**)), it might seem reasonable to specify GWA inference for MAP-SSVS in a similar manner; i.e, using the E-step values of τ*_j_* at convergence as estimates of PPA. However, we noted that these E-step values uniformly drifted either towards 0 or 1 such that there were never any intermediate estimates of PPA. A comparison of PPA based on *τ_j_* for Prob(*τ_j_* =1|**y**) for MCMC-SSVS versus the E-step values of *τ*at convergence on the MSUPRP data is provided is given in Panel A of Figure S3 in Supplementary File S2. Also, recall that the MAP-procedure is sensitive to starting values and that starting values for MAP-SSVS were based on RRBLUP as this might be a pragmatic and reasonable strategy in most cases. If we had based starting values on, say, their MCMC-SSVS posterior means, one would notice a different assortment of converged Estep values of *τ*compared to what we observed with RRBLUP starting values as we demonstrate with the MSUPRP data in Panel B of Figure S3.

Recall that for MAP-SSVS, we based starting values for the SNP specific PPA on estimated local false discovery rates (lFDR) using the R package ashr since there is presumably a close relationship between them; i.e., PPA ≈ 1- lFDR (Stephens 2017). This procedure converts *P*-values to lFDR such that we based lFDR determinations from the *P*-values computed under EMMAX. This begged the question as to whether PPA could be simply based on lFDR processing of EMMAX *P*-values. However, upon comparing 1-lFDR estimated from the EMMAX P-values to PPA estimated using MCMC-SSVS of the MSUPRP data, it appeared that there was not generally very good agreement between the two sets of PPA estimates except for the some near-zero PPA and the largest PPA estimated using both procedures (Figure S4 in Supplementary File S2).

We also wondered if the strategy for computing window-based PPA could be simplified further from that presented in Fernando *et al*. (2014) and used in this paper (i.e., Equation [19]) to that suggested by Moser *et al*. (2015) who simply summed SNP specific PPA (i.e., based on Equation [15]) within a window to determine the window-based PPA. One should anticipate that the approach of Moser *et al*. (2015) should lead to higher estimated PPA. We compared the two PPA determination approaches for pAUC05 in the simulation study and noted that there was significant interaction between PPA determination approach with γ and *n_qtl_* but no significant interaction involving window size; hence we compared the two strategies within each value of γ and *n_qtl_* averaged across window length (Table S7 in Supplementary File S2). The only significant difference in pAUC05 occurred with γ=3 and *n_qtl_* =300 for which the approach of Fernando *et al*. (2014) lead to a higher pAUC05. Nevertheless, since point estimates of pAUC05 were always larger using the approach from Fernando et al. (2014), we would recommend their approach from Equation [19] for the determination of windows based PPA.

We did not estimate *ν_τ_* using either the procedures outlined in Yang *et al*. (2015) for MCMC-BayesA or provided in Chen and Tempelman (2015) for MAP-BayesA primarly because of the extremely poor MCMC mixing for sampling this hyperparameter and its poor convergence in MAP-BayesA. A typical specification for *ν_τ_* in BayesA is 4 or 5 (Colombani *et al*. 2013; Perez and de los Campos 2014). The specification of *ν_τ_* = 2.5 that we chose for this paper was based in part on results from Yang *et al*. (2015) and Nadaf *et al*. (2012) who determined that lower specifications of v (i.e., heavier tails) could lead to higher genomic prediction accuracies when using Bayes A. To assess this further, we compared MCMC-BayesA using *ν_τ_* = 2.5 versus *ν_τ_* = 5 for pAUC05 based on the `BALD` derived adaptive window inference. In general, the use of *ν_τ_* = 2.5 yielded a higher mean pAUC05 than *ν_τ_* = 5 except for a non-significant difference at *n_qtl_*=*300* (Table B4). For large scale empirical analyses whereby hyperparameter inference seems daunting, researchers should consider conducting analyses based on a finite number of hyperparameter specifications, choosing those specifications that lead to the best cross-validation prediction accuracy. Similar arguments could be made for choosing the key hyperparameters in other Bayesian regression models. It is worth noting that even we ran our MCMC algorithm for 1 million iterations, the mixing of the MCMC chain was still rather poor as it pertained to inference on other hyperparameters. For example, for MCMC-BayesA, the effective sample size (ESS) for 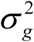 was estimated to be 66.33 whereas for SSVS, the ESS was 61.03 for 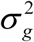 and 53.48 for *π*_τ_.

It should be apparent that given that MCMC-SSVS is a natural variable selection model, it might be favored over MCMC-BayesA which is not a natural variable selection model. Our strategy for computing the proportion of genetic variance explained by each window and determining the posterior probability that that percentage exceeds an arbitrary threshold (1% in our analyses) is based on the strategy presented by Fernando and Garrick (2013). The flexibility of MCMC modeling allows posterior probabilities (i.e., PPA) of this nature to be computed. However, one should be wary of the impact of the threshold since it obviously should depend upon marker density. That is, if the threshold is set to be too high, then sensitivity is lost. Based on the results from both simulation study and real data analysis, we demonstrated that random effects modeling can also be powerful tool for GWA as long as the suitable priors, i.e., in our case sparser priors, are used. Other variable selection implementations popularized in WGP including BayesB (Meuwissen *et al*. 2001) or BayesR (Erbe *et al*. 2012; Moser *et al*. 2015) could be considered as well.

Our MSUPRP application was interesting in that we discovered that SNP markers in two different blocks can be in high LD even when they’re not adjacent to each other. However, we would quickly note that these strange LD patterns may be due to genome assembly errors in the pig genome (Groenen 2016) with particular issues having been identified in the Chromosome 6 region (Warr *et al*. 2015) which contained the strongest associations in our study. This may somewhat complicate strategies for single SNP specific or even window-based inference. We also recognize that there is a movement towards the use of multi-SNP haplotype modeling which may improve GWA performance (Cuyabano *et al*. 2014). Our adaptive window based strategy seems to improve the performance of GWA relative to single SNP or fixed window length inference although, conceivably, there may be other better ways to group SNPs. With marker densities well beyond 50K, the adaptive window strategy might not be viable since it requires the computation and storage of matrix of LD r^2^ values between every SNP marker within a chromosome before clustering analyses can be used to partition the genome into windows. Fernando *et al*. (2014) also suggested that PPA based on Bayesian GWA analyses similar to our MCMC-SSVS be based on whether non-zero associations were found not only in that marker’s resident window but also in either of the two flanking windows. Their strategy was based on fixed window sizes such that it may be worthwhile to consider their flanking strategy in the context of adaptively chosen window sizes. We conjecture that if LD structure is appropriately used to partition the genome, the use of such flanking windows might not be necessary; however, this should be a topic for future research. It is also important to note that the comparisons in this paper are context specific in terms of the genomic LD relationships germane to a F2 cross in pigs. This naturally leads to a higher pairwise LD between adjacent SNP markers than what might be found in outbreeding populations, and most notably humans. This naturally would change the relative comparisons between single SNP versus windows based inferences as well as the relative number and sizes of adaptively chosen windows based on LD relationships. Hence future investigation of our approaches in other populations is strongly warranted.

In summary, we found Bayesian variable selection to be a promising strategy for GWA when combined with window based inference. Nevertheless, it seems prudent that window selection be carefully chosen using rules based on LD information rather than predetermined constant physical window lengths (in Mb) for genomic regions. Also, recently proposed analytical approaches for Bayesian regression models should be discouraged for GWA studies.

## ACKNOWLEDGMENTS

We are indebted to the group of Dr. Cathy Ernst for making this data available. We are also grateful to Jose-Luis Gualdron Duarte and Yeni Bernal Rubio for their assistance in preparing the data. This project was supported by Agriculture and Food Research Initiative Competitive Grants no. 2011-77015-30338 and 2004-35604-14580 from the United States Department of Agriculture National Institute of Food and Agriculture.

